# Investigating the significance of iron levels in influencing megakaryocytic commitment in megakaryocyte-erythroid progenitors

**DOI:** 10.64898/2026.08.11.743878

**Authors:** Ranita De, Leo Stephen, Vikram Mathews, Sajitha Lulu, Akshayata Naidu, Blessy Kiruba, Paweł Lipiński, Rafał Starzyński, Eunice Sindhuvi Edison

## Abstract

**Aim:** The present study investigated the significance of iron in regulating megakaryopoiesis, by a diet-based intervention in an *in-vivo* model.

**Methods:** Male C57BL/6 mice, aged 4-5 weeks were fed on varying iron diets. Following sacrifice, blood samples collected in EDTA tubes were used to analyse haematological parameters, and iron content of liver and spleen was assessed by biochemical analyses. Megakaryocyte-erythroid progenitors (MEPs) were isolated from bone marrow by magnetic bead-based selection. RNA isolated from bone marrow cells and MEPs were used for gene expression analyses, and RNA Sequencing to identify differentially expressed genes (DEGs) and associated pathways.

**Results:** Mice fed on an iron-deficient diet had reduced hepatic iron content after 5 weeks (p < 0.01), while both the hepatic and spleen iron content increased after 3 weeks in mice on an iron-rich diet (p < 0.05) and developed iron overloading. Hb and RBC counts increased (p < 0.05) in iron-rich mice and decreased in iron-deficient mice (p < 0.05), which also showed elevated platelet counts (p < 0.01). This may be explained by increased expression of Gata1, Tal1 (p < 0.01) Mds1 and Pdpk1 (p < 0.05) in bone marrow cells from iron-deficient mice. MEPs isolated from these mice showed elevated expression of genes associated with megakaryocytic differentiation, platelet functions, and genes encoding TGF-βR1 and Smad 2,3 and 4.

**Conclusions:** Iron deficiency may activate TGF-β signalling and downstream Smad-mediated transcriptional programs within MEPs. This may promote a shift in lineage commitment towards megakaryopoiesis through elevated expression of megakaryopoiesis related genes.

## Introduction

Platelets or thrombocytes are central players of hemostasis, which are conserved in all vertebrates. Their multifunctional nature renders them indispensable for the initiation of coagulation cascades at the surface of damaged blood vessels, apart from playing a pivotal role in disease pathophysiology [1]. Normal platelet counts in healthy adults’ ranges from 150 – 450 × 10^9^/L. An increase in their counts beyond 450× 10^9^/L is known as “thrombocytosis” and “thrombocytopenia” refers to a drop in their counts below 150 ×10^9^ /L. More than 85% of cases of thrombocytosis are associated with secondary causes such as infections, iron deficiency anemia (IDA), post-splenectomy cases, trauma, non-infectious inflammation or blood loss [2]. A close relationship has been documented between body iron stores and platelet counts. IDA has been identified as a common causative factor of reactive thrombocytosis [3] and less frequently thrombocytopenia [4].

The significance of an individual’s iron status towards regulation of platelet counts is not well elucidated. Several *in vivo* and *in vitro* studies have been conducted to understand probable mechanisms through which iron levels may influence megakaryopoiesis. Platelets obtained from iron deficient rats displayed altered characteristics including increased counts, size and aggregation. These changes were unlikely to be mediated by thrombopoietic cytokines such as IL-6, IL-11 and thrombopoietin (TPO), as iron deficient rats and controls had similar levels of these cytokines [5]. A search for potential target genes regulating megakaryopoiesis in iron deficiency indicated an up regulation of Hypoxia-inducible factor 1, α subunit (HIF2α) and its downstream target, vascular endothelial growth factor (VEGF-A) in erythroleukemic cell lines, propagated in presence of iron deficient conditions [6]. Recently Liu et al. have reported that mice affected with IDA have impaired platelet functions, while high iron concentrations activated platelets due to enhanced collagen-induced calcium mobilization. Elevated iron levels also promoted integrin αIIbβ3 signaling and ferroptosis in platelets, thereby increasing their activation [7].

According to the classical model of haematopoiesis, megakaryocytes or platelet precursors arise from megakaryocyte-erythroid progenitors (MEPs), which are derived from common myeloid progenitors (CMPs, with myeloid, erythroid and megakaryocytic potential) [8]. However, this model of the origin of different haematopoietic lineages, especially that of megakaryocytes, has been challenged in recent years. Although different surface markers have been utilised to define MEPs [9], megakaryocytes and erythroid cells probably share a common progenitor as they are the first terminally differentiated blood cells to be produced during embryonic haematopoiesis [10]. Ferrucio *et al.* carried out an elegant study in mice where transmembrane serine protease 6 (Tmprss6) was knocked out [11]. Lack of functional Matriptase-2 (MT2) encoded by Tmprss6, results in high levels of the iron-regulatory hormone hepcidin, which prevents iron absorption and leads to iron-refractory iron-deficiency anemia [12]. Tmprss6-/-mice exhibited thrombocytosis along with IDA. MEPs isolated from these mice were biased towards the megakaryocyte lineage, and showed an increased expression of genes involved in metabolic, extracellular signal-regulated kinase (ERK) pathways compared to wild type MEPs [11].

In the present study we investigated the effect of varying iron levels on megakaryopoiesis by maintaining C57BL/6 mice on control, iron-deficient and iron-rich diets. The significance of a diet-based intervention on iron status was determined by biochemical analyses of hepatic iron stores and analyses of haematological parameters. RNA isolated from bone marrow cells was used to study the expression of megakaryocytic lineage-specific genes. This model was further refined to investigate changes in haematological parameters and iron stores in mice fed on varying iron diets for different time intervals. RNA isolated from a pure population of MEPs was subjected to RNA Sequencing and gene expression analyses to identify differentially expressed genes (DEGs) and associated pathways. The present study investigates the mechanistic significance of iron in regulating the transcriptional switch towards megakaryopoiesis in MEPs.

## Materials and methods

### Mice

Male C57BL/6 mice, aged 4-5 weeks and weighing 20-25 g were used. They were maintained at 22±3°C and 55±15% relative humidity with a 12-hour light-dark cycle. Food and water were provided ad libitum. Two sets of mice were used. Studies carried out at the Institute of Genetics & Animal Biotechnology of the Polish Academy of Sciences, were approved by the 2^nd^ Local Ethical Commission at the Warsaw University of Life Sciences, Poland (Permission No-WAW2/054/2020). The second set of animal studies were approved by the Institutional Animal Ethics Committee (IAEC) of Christian Medical College, Vellore (IAEC No-5/2018). Both sets of mice were fed an iron-deficient diet (3 ppm iron), an iron-rich diet (20,000 ppm iron) and a control diet (40 ppm iron), Research diets, Inc. NJ, USA) for different time durations. The mice models developed are summarized in Supplementary Figure 1.

### Model I

A preliminary model developed to investigate the effect of varying iron levels on megakaryopoiesis in an *in vivo* context. Male C57BL/6 mice were subjected to a diet-based intervention, and haematological parameters and hepatic iron content were assessed. Bone marrow cells were isolated from mice fed on varying iron diets to analyse expression of megakaryocytic lineage-specific genes.

### **a)** Measurement of haematological parameters

Following propagation on iron-deficient diet (n=3), control diet (n=3) for 5 weeks and an iron-rich diet (n=3) for 3 weeks, mice were sacrificed by cardiac puncture. Blood samples were collected in EDTA tubes and processed within 4 hours to analyse haematological parameters using the ADVIA 120 haematology analyser.

### **b)** Assessment of hepatic iron content

Liver samples from mice were fixed in Bouin’s solution for 24 hours. They were then stored in 70% ethanol at room temperature. Haem and non-haem iron content was analysed by the Accustain iron deposition kit.

### **c)** Isolation of cells from mice bone marrow

Bones from the tibia, femur, and clavicles were scraped and rinsed with sterile PBS. Bone marrow cells were flushed and collected in ice-cold sterile PBS. These cells were pelted down by gentle centrifugation at 1200 rpm for 5 minutes and resuspended in Trizol and stored at-80°C for subsequent RNA extraction.

### **d)** Gene expression studies

Total RNA was extracted from bone marrow cells using Trizol reagent (Invitrogen, CA, USA) according to standard protocol. The RNA pellet isolated was air-dried, suspended in an adequate volume of RNase-free water, and stored at-80°C for further use. The concentration and purity of RNA were assessed using a nanodrop microvolume UV-Vis spectrophotometer (Thermo Scientific, USA). RNA isolated was used for cDNA synthesis by the RT2 First Strand kit (Qiagen) as per the manufacturer’s protocol. 500 ng of RNA was utilised for cDNA synthesis. The latter was used for real-time quantitative PCR in a Light Cycler U96 system (Roche Diagnostics). Amplified products were detected using SYBR Green (Takyon). Plate-to-plate variation was controlled by normalising gene expression with respect to the housekeeping gene i.e. Gapdh, by using the 2^-ΔΔCt^ method.

### Model II

In the previous model, iron content and expression of megakaryocytic genes were assessed at only one time point in mice fed on varying iron diets. We did not investigate how these factors may vary as iron deficiency/overloading develop. Total bone marrow cells were used to study the expression of megakaryocytic lineage-specific genes. This does not paint a true picture of the significance of iron in regulating megakaryopoiesis, due to the heterogeneous progenitor cell populations found in the marrow. These shortcomings were addressed in a refined mouse model.

### **a)** Measurement of haematological parameters

Mice fed on an iron-deficient diet (n=3), an iron-rich diet (n=3), and a control diet (n=3) for different time durations (Supplementary Fig 1), were sacrificed by cardiac puncture. Blood samples in EDTA tubes were processed to analyse haematological parameters using the Sysmex XN – 1000V (Sysmex Corporation, Kobe, Japan) haematology analyser.

### **b)** Assessment of iron content of principal iron storage organs

Tissue samples collected from liver and spleen were rinsed with PBS, and stored in nitric acid and hydrogen peroxide (3:1 ratio). Their iron content was quantified by atomic absorption spectroscopy. Histochemical assessment of hepatic iron content was performed by rinsing liver tissues from mice fed on different iron diets in PBS (1X), followed by storage in a fixative solution (10% formalin). These were then subjected to Perl’s staining.

### **c)** Isolation of MEPs from mice bone marrow cells

#### i) Isolation of Lin^-^ ckit^+^ Sca1^-^ cells from mice bone marrow

To isolate megakaryocytic progenitor cells, bone marrow cells were collected in ice-cold sterile PBS and processed further. Briefly, RBCs were removed from these cells by incubation in the presence of ACK lysis buffer for 5 minutes at room temperature. The cell pellet obtained was washed with PBS and resuspended in a conditioning medium (PBS containing 2% FBS and 0.5 mM EDTA) for isolation of Lineage^-^ cells using the EasySep™ Mouse Haematopoietic Progenitor Cell Isolation kit (Stem Cell Technologies) as per the manufacturer’s protocol. The isolated Lin^-^ckit^+^ Sca1^-^ cells were resuspended in the conditioning medium for subsequent processing.

#### **ii)** Isolation of CD34^-^ cells from Lin^-^ ckit^+^ Sca1^-^ cells

The Lin^-^ckit^+^Sca1^-^ cells isolated from bone marrow were resuspended in the required volume of conditioning medium. These cells were then subjected to CD34 depletion by negative selection by using the EasySep™ mouse PE positive selection kit (Stem Cell Technologies) as per the manufacturer’s protocol. A fraction of the isolated MEP cells (5 × 10^5^ cells) was utilised for flow cytometric analyses of the expression of surface markers.

### **d)** Flow cytometry

MEP cells were pelleted down by gentle centrifugation, washed with PBS supplemented with 0.5% NaN3 (Sigma-Aldrich) and 0.1% FBS, and resuspended in PBS. Following this, they were stained for 20 minutes with anti-mouse pre-conjugated antibodies i.e. PE-labelled CD117 (c-kit), FITC-labelled Ter119, PE-labelled CD34 and APC-labelled Ly6A/E (Sca 1) (BD Pharmingen, San Diego, CA, USA). Excess antibody was removed by washing cells with PBS and acquired on a Navios flow cytometer (Beckman Coulter). Viable cells were gated by staining with Per-CP labelled 7-AAD (BD Pharmingen, San Diego, CA, USA). The data analysis was performed using the Kaluza software (Beckman Coulter).

### **e)** Molecular analyses

Total RNA was extracted from isolated MEP cells using Trizol reagent (Invitrogen, CA, USA) according to standard protocol. The concentration and purity of RNA pellet was assessed as previously described. 500 ng of isolated RNA isolated was used for cDNA synthesis as described beforeExpression of target genes was quantified using the 7500 QPCR System (Applied Biosystems). The primer sequences of megakaryocytic lineage specific transcription factors and certain novel genes, whose functions were associated with megakaryocyte development, platelet activation, and signalling are listed in Supplementary table 1.

### **f)** RNA Sequencing

RNA extracted from MEP cells, isolated from bone marrow cells of mice maintained on different iron diets was subjected to RNA Sequencing (Illumina Platform). The raw reads obtained were filtered using Trimmomatic for quality scores and adapters. Filtered reads were aligned to the Mus musculus genome using a splice-aware aligner like HISAT2 to quantify reads mapped to each transcript. The alignment percentage of reads was in the range of 96.09-98.57%.

The total number of uniquely mapped reads was counted using feature counts. The uniquely mapped reads were then subjected to differential gene expression using Deseq2 software (Supplementary figure 1). The obtained DEGs were filtered based on p values < 0.05. The filtered genes were put in the Network Analyst and Reactome database to find their relevant transcription factors and pathways, respectively.

### **g)** Statistical analyses

Quantitative variables were reported using mean ± standard deviation (SD) or median and interquartile range (IQR), according to the data distribution characteristics. For qualitative variables, numbers and percentages were used. Differences in means from non-repeated measures were assessed using ANOVA or Kruskal-Wallis test, for parametric and non-parametric data respectively (Graph Pad Prism 8.0.1). For all tests, a p-value of < 0.05 was considered as being statistically significant.

## Results

### 1. Model I - Iron-depleted diet affects hepatic iron content and haematological parameters in mice

Our first goal was to establish a robust mice model to study effects of iron deficiency and overloading by a diet-based intervention. Mice fed on an iron-deficient diet for 5 weeks had significantly reduced hepatic haem and non-haem iron content, compared to those fed on an iron-rich diet (p < 0.001). Being fed on an iron-deficient diet also resulted in significantly reduced non-haem iron than mice exposed to a control diet (p < 0.05) (Figure 1E).

**Figure 1.**
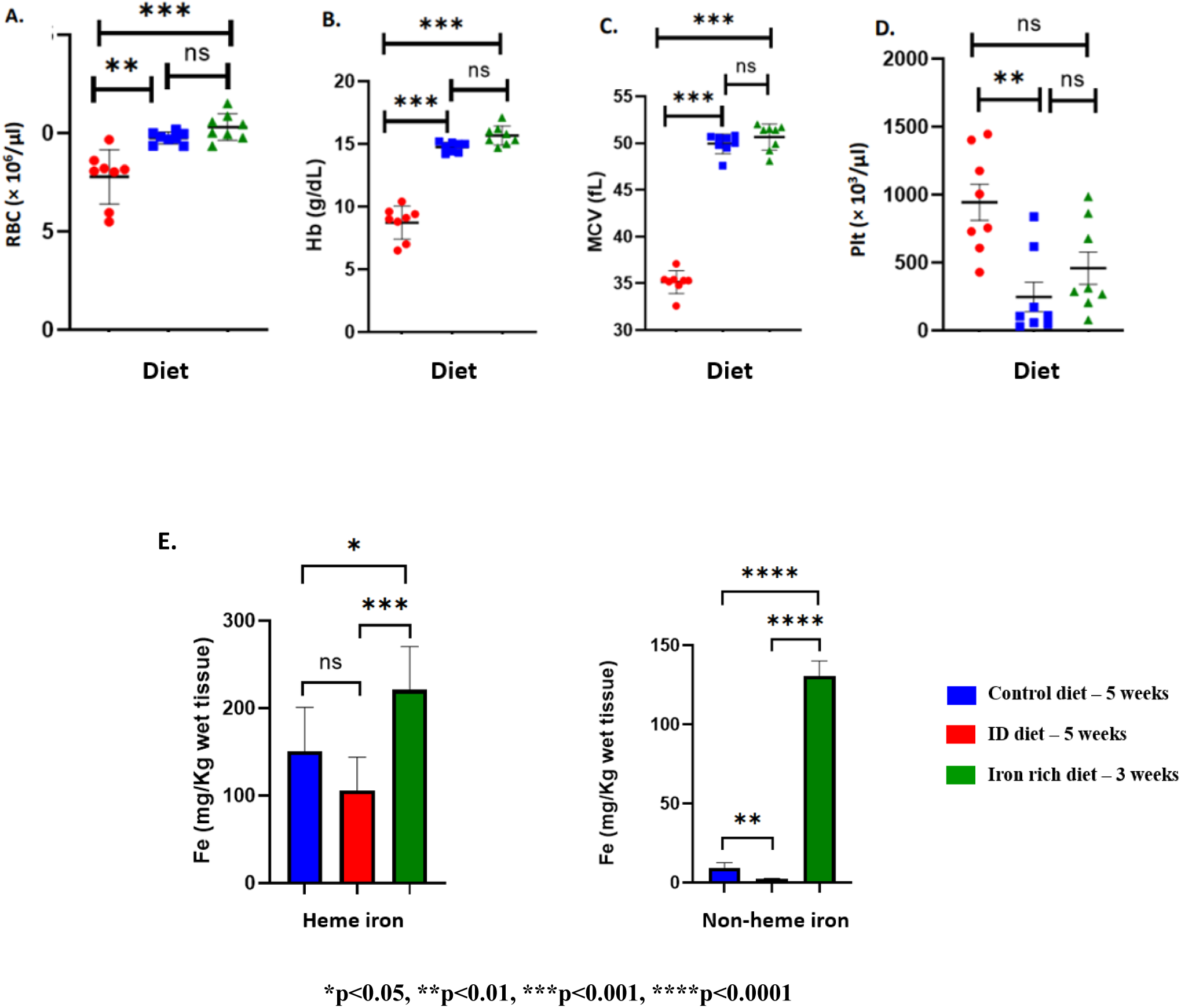
Iron-deficient diet induces iron deficiency and anaemia in mice –. Mice maintained on an iron-deficient diet for 5 weeks have significantly reduced RBC counts (p<0.01), Hb (p<0.001) and MCV (P<0.001), compared to control mice (A-C), but increased platelet counts (p<0.01) (D). Iron-deficient diet significantly decreases hepatic heme (p<0.001) and non-heme iron (p<0.0001) (E), compared to mice fed on an iron-rich diet for 3 weeks. n=3 for each group.

These iron-deficient mice had significantly reduced RBC counts, Hb and MCV, indicative of anaemia development. Interestingly, these mice had significantly increased platelet counts (880 × 10^3^/µl, 429-1446 × 10^3^/µl) compared to control mice (108 × 10^3^/µl, 30-838 × 10^3^/µl) (p < 0.01) (Figure 1 A-D).

#### 1.1. Expression of megakaryocytic lineage specific genes increases in bone marrow cells from iron-deficient mice

Next, we analysed the effect of iron deficiency on megakaryopoiesis. RNA isolated from total bone marrow cells of mice fed on varied iron diets were used for analysing expression of megakaryocytic lineage-specific genes, which were identified after literature screening.

Gata1 was significantly increased (∼ 3-fold) (p < 0.01) in mice fed on an iron-deficient diet for 5 weeks. Tal1 was also significantly upregulated (∼3-fold) (p<0.01), which forms a complex with other transcription factors and induces megakaryopoiesis. In an interesting observation, iron deficient mice showed significantly increased upregulation of Mds and Evi1 complex locus (Mds1) (∼ 2-fold) (p < 0.05), and 3-phosphoinositide dependent protein kinase 1 (Pdpk1) (1.8-fold) (p < 0.05) (Figure 2). These genes have been recently reported to regulate megakaryocytic differentiation and proplatelet formation stages.

**Figure 2.**
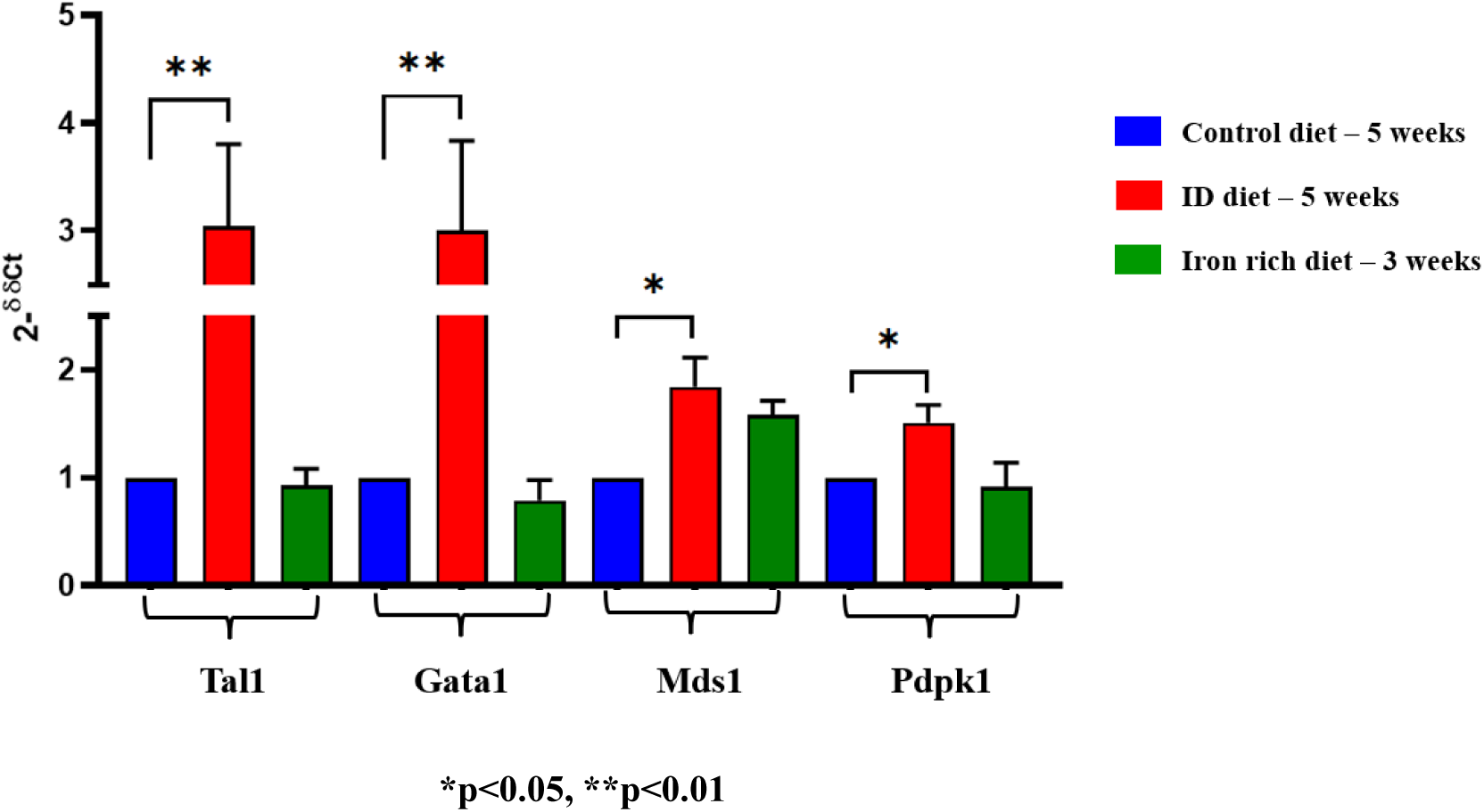
Expression of megakaryocytic lineage-specific genes & transcription factors are upregulated in bone marrow cells isolated from iron-deficient mice –. Megakaryocytic transcription factors Tal1 and Gata1 showed significantly increased expression (p<0.01) in iron-deficient mice, compared to control mice. These mice also showed significantly increased expression of Mds1 and Pdpk1(p<0.05), which have been recently associated with regulating megakaryocytic differentiation and proplatelet formation. Reference gene – Gapdh, n=3 for each group.

**Figure 3.**
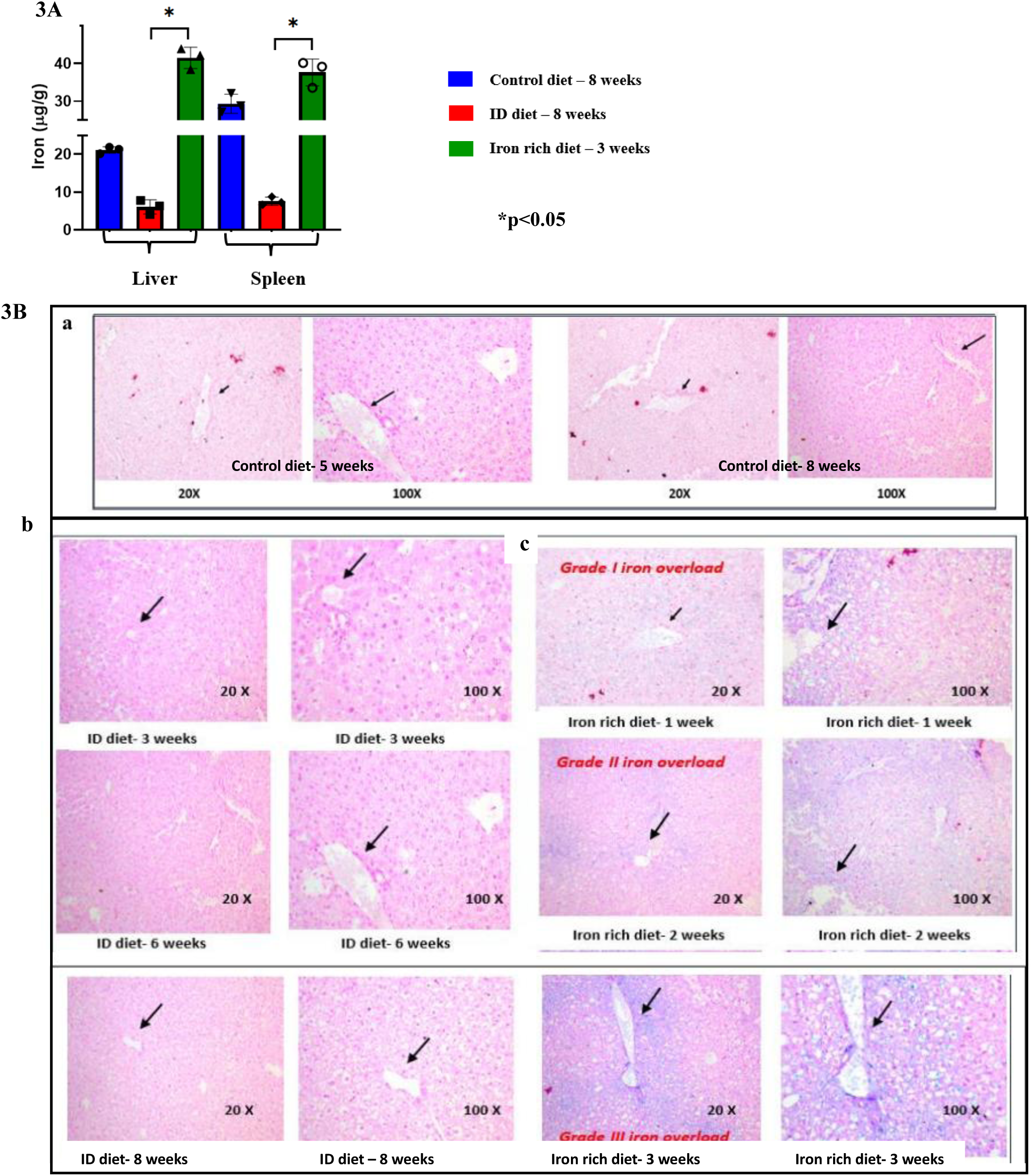
Mice on an iron-deficient diet have decreased hepatic, spleen iron content and develop iron overloading on an iron-rich diet –. Exposure to an iron-deficient diet significantly decreases iron content of principal iron storage organs (p<0.05), compared to mice fed on an iron-rich diet (A). Assessment of hepatic iron content by Perl’s staining indicates no stainable iron granules were observed in liver sections extracted from control mice (3B, a). Similar results were observed in mice fed on an iron-deficient diet (3B, b). Zonal distribution of iron granules was observed in mice maintained on an iron-rich diet; iron overloading progressed from Grade I to III after 3 weeks (3B, c). Above images depict liver parenchyma with portal tract (arrow) and iron granules (purple). n=3 per group.

### 2. Establishment of a refined mouse model maintained on varying iron diets for different time durations

To test how iron content and haematological parameters may vary during the development of iron deficiency mice fed on iron-deficient diet (n=3) were sacrificed after 3 weeks, 6 weeks and 8 weeks. Mice maintained on an iron-rich diet (n=3) were sacrificed after 1 week, 2 weeks and 3 weeks, while those on a control diet (n=3) were sacrificed after 5 weeks and 8 weeks. In the previous model, we observed an increased expression of megakaryocytic lineage specific genes and transcription factors in bone marrow cells isolated from iron-deficient mice. However, to gain an in-depth understanding of the significance of iron in influencing megakaryopoiesis, a pure population of megakaryocyte-erythroid progenitors (MEPs) were isolated from bone marrow cells. Gene expression profile of these MEPs were analyzed in mice that had developed iron deficiency and overloading.

#### 2.1. Iron-deficient mice gradually develop anaemia and thrombocytosis

After being fed on an iron-deficient diet for a week, RBC counts were 7.96 ± 0.15 × 10^6^ /µl, Hb was 11.5 ± 0.87 g/dl and MCV was 46.36 ± 0.55 fl. These declined significantly after 3 weeks, wherein RBC counts were 6.52 ± 0.51 × 10^6^ / µl (p < 0.05), Hb was 8.9 ± 1.01 g/dl (p < 0.05) and MCV was 45.3 ± 0.57 fl. No significant difference in these parameters was observed in control mice over the same time interval. However, these parameters displayed an increasing trend in mice, fed on an iron-rich diet. In these mice, RBC, Hb and MCV were 8.24 ± 0.26 × 10^6^ / µl, 12 ± 0.1 g/dl and 45.6 ± 0.55 fl, respectively after 1 week. They significantly increased to 10.61 ± 0.71× 10^6^ / µl (p < 0.05), 17.2 ± 1.1 g/dl (p < 0.01), after 3 weeks. Platelet counts significantly increased from 323 × 10^3^ / µl (212-342) after 1 week, to 622 × 10^3^ / µl (593-709) (p < 0.01) after 3 weeks on an iron-deficient diet (Figure 4).

**Figure 4.**
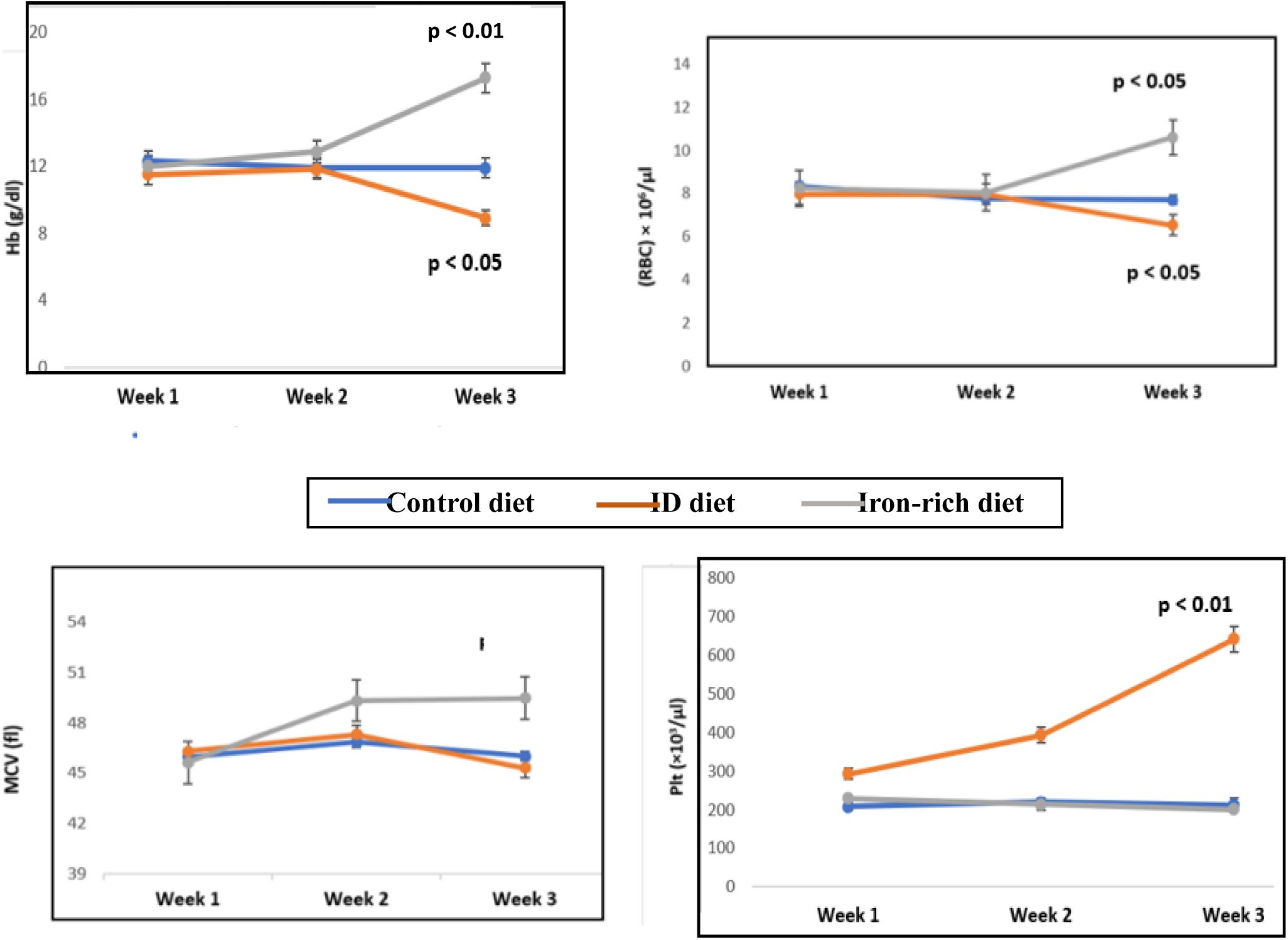
Iron-deficient mice gradually develop anaemia and thrombocytosis. -RBC counts and Hb decrease significantly after 3 weeks (p<0.05), compared to the first week in mice maintained on an iron-deficient diet. Both RBC counts (p<0.05) and Hb (p<0.01) increase significantly after 3 weeks in mice fed on an iron-rich diet. Platelet counts increase significantly (p<0.01) in iron-deficient mice during this time interval. n=3 per group.

#### 2.2. Iron-deficient diet affects iron content of iron-storage organs and mice develop iron overloading on an iron-rich diet

The iron content of the liver and spleen in mice fed on varying iron diets was analysed by AAS. After being fed on an iron-rich diet for 3 weeks, the hepatic and spleen iron content significantly increased (p < 0.05). On the contrary, the iron content of both these tissues decreased in mice fed on an iron-deficient diet for 8 weeks, compared to those fed on a control diet for the same time. These differences were not significant. Changes in tissue iron content have been depicted in Figure 3A.

Next, we performed histochemical assessment of hepatic iron content by Perl’s staining. No stainable iron granules were found in liver parenchyma, in mice fed on iron-deficient (Figure 3B b) and control diets (Figure 3B a) for different time durations. However, mice developed Grade I iron-overloading after just 1 week on an iron-rich diet, which progressed to Grade III after 3 weeks (Figure 3B c).

#### 2.3. Genes regulating megakaryopoiesis and platelet homeostasis and functions are up regulated in MEPs isolated from iron-deficient mice

We wanted to investigate the significance of iron levels in regulating megakaryocytic commitment in MEPs. Thus, RNA extracted from MEPs (Lin^-^ckit^+^ Sca1^-^ CD34^-^), which were isolated from bone marrow cells of mice fed on different iron diets were subjected to RNA sequencing (Figure 5A). We found different genes involved in regulating megakaryocytic development, platelet homeostasis and functions were upregulated in iron-deficient MEPs. Conversely, various genes involved in platelet activation and homeostasis were downregulated in MEPs isolated from mice fed on an iron-rich diet (Table 1).

**Figure 5.**
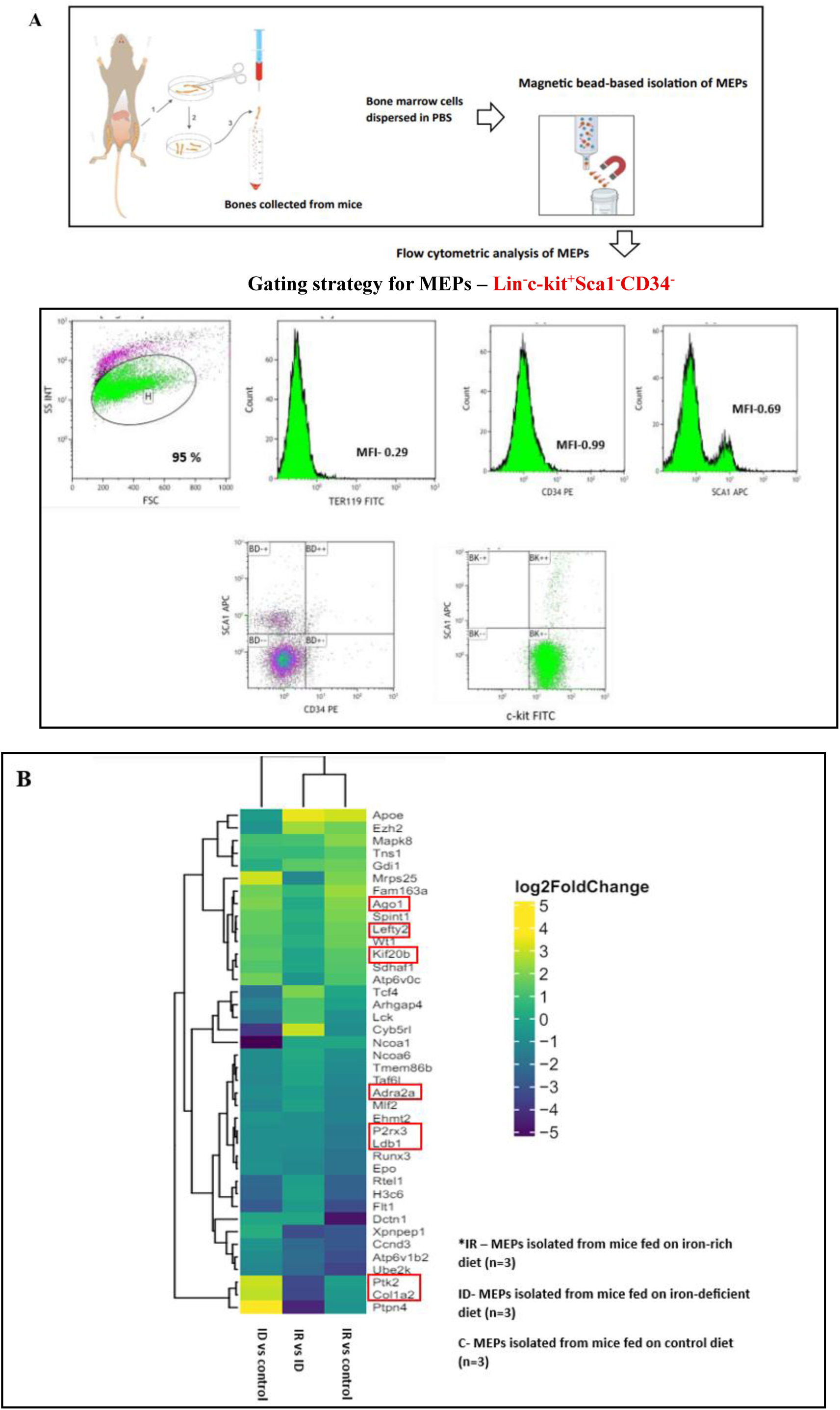
**Genes regulating megakaryopoiesis & platelet functions are up-regulated in iron-deficient MEPS**-Protocol for isolation of MEPs (A). Heatmap of differentially expressed genes in MEPs isolated from control, iron-deficient and iron-rich mice (B). Genes associated with megakaryopoiesis & platelet functions are highlighted in red. n=3 per group

**Table 1.** Summary of DEGs and megakaryopoiesis associated pathways in MEPs isolated from iron-deficient and iron-overloaded mice.

| Gene | Full name | Associated function identified |
| --- | --- | --- |
| Lefty2 | l-right determination factor 2 | telet degranulation, signalling, regulates resp<br>levated cytosolic calcium. |
| Ago1 | gonaute RISC component 1 | mulates RUNX1 regulated genes, involve<br>gakaryocytic differentiation. |
| Kif20b | esin family member 20b | gulates factors involved in megakaryo<br>elopment & platelet production. |

B
| Gene | Full name | Associated function identified |
| --- | --- | --- |
| Gnb2 | G protein subunit beta 2 | gulates platelet homeostasis. |
| P2rx3 | Purinergic receptor P2X3 | gulates platelet homeostasis. |
| Adra2a | Adrenoceptor alpha 2a | gulates platelet aggregation, activation & signalling |
| Ldb1 | LIM domain binding 1 | gulates platelet calcium homeostasis, med<br>gakaryocytic differentiation as a RUNX1 target gen |

- (A) DEGs up-regulated in MEPs isolated from mice fed on an iron-deficient diet for 8 weeks (n=3). - (B) DEGS downregulated in MEPs isolated from mice fed on an iron-rich diet for 3 weeks (n=3).

The heat map depicting differentially upregulated and downregulated genes in MEPs, isolated from mice fed on control, iron-deficient and iron-rich diets is depicted in Figure 5B. Genes associated with megakaryocytic development, platelet homeostasis and functions have been highlighted.

#### 2.4 Gene expression analyses of DEGs identified by RNA sequencing

DEGs from different mice groups were filtered based on p values < 0.05, relevant transcription factors and pathways were identified from the Network Analyst and Reactome database, respectively. Genes associated with megakaryocytic development, differentiation, platelet homeostasis and functions were shortlisted, and their expression was validated by real time PCR studies.

Iron-deficient MEPs (n=3) showed significantly increased upregulation of Lefty2 (7.5-fold) (p < 0.01), Gnb2 (> 6-fold) (p < 0.01) and P2rx3 (> 5-fold) (p < 0.01). Apart from these genes, Adra2a (5-fold), Ldb1 (7-fold) and Ago1 (3-fold) were also increased in this cohort, compared to control MEPs. These changes were not significant. Conversely, Lefty2 (∼ 1-fold) (p < 0.01), Gnb2 (∼ 1-fold) (p < 0.01) and P2rx3 (0.6-fold) (p < 0.01) were significantly reduced in iron-rich MEPs, compared to iron-deficient MEPs. MEPs from iron-rich mice also showed significantly reduced expression of Ldb1 (0.87-fold) and Kif20b (0.38-fold) (p < 0.05) (Figure 6).

**Figure 6.**
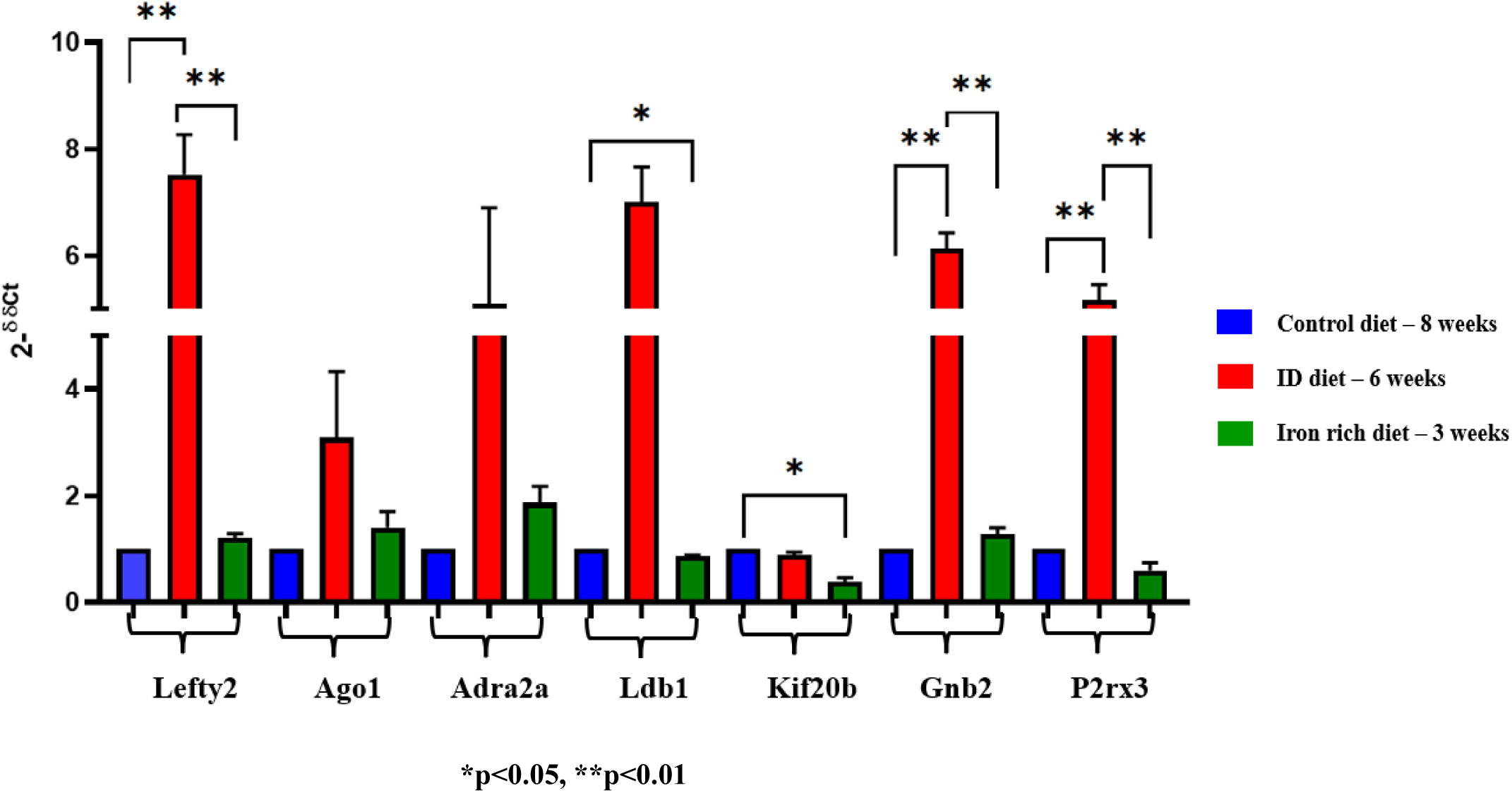
Genes regulating megakaryocytic development & platelet functions show increased expression in iron-deficient MEPs. – Lefty2, associated with platelet degranulation, signalling and Gnb2, P2rx3 associated with regulation of platelet homeostasis showed significantly increased expression (p<0.01) in iron-deficient MEPs, compared to controls. These genes were significantly down regulated in iron-rich MEPs (p<0.01). Genes linked with megakaryocytic differentiation and platelet production such as Kif20b and Ldb1 showed significantly reduced expression in iron-rich MEPs (p<0.05), compared to controls. n=3 per group.

##### 2.4.1. Genes regulating megakaryocytic development and platelet functions showed increased expression over time in iron-deficient MEPs and were downregulated in iron-rich MEPs

We assessed temporal changes in gene expression in iron-deficient and iron-rich MEPs. Lefty2, Gnb2 and P2rx3 showed low expression i.e. 0.65-fold, 0.1-fold, and 0.43-fold, respectively, in iron-deficient MEPs after 3 weeks. They significantly increased to 7.5-fold, ∼ 6-fold and ∼ 5-fold, respectively, after 6 weeks (p < 0.01). Other genes, including Ago1 (3-fold), Adra2a (5-fold) and Ldb1 (7-fold), were also upregulated in MEPs from iron-deficient mice after 6 weeks, in comparison to 3 weeks. These differences were not significant (Figure 7A).

**Figure 7.**
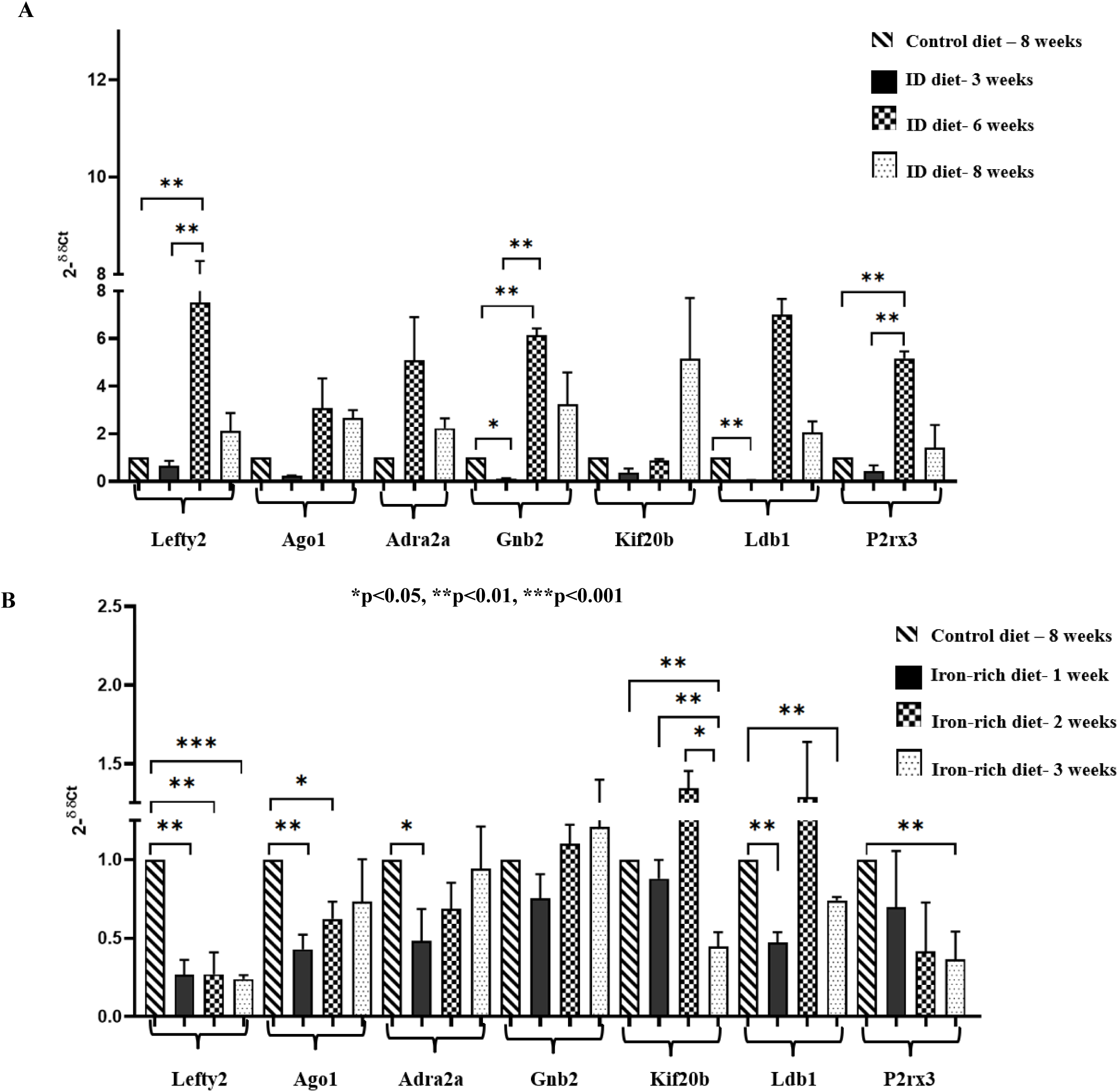
Genes regulating megakaryopoiesis and platelet functions showed temporal increase in expression in iron-deficient MEPs and were downregulated in iron-rich MEPs –. Expression of Lefty2, Gnb2 and P2rx3 increased significantly from week 3 to 6 in iron-deficient MEPs (p<0.01) (A). Lefty2, Ago1, Adra2a and Ldb1 showed significantly reduced expression (p<0.01) after week 1 in iron-rich MEPs compared to controls. Expression of Ldb1, P2rx3 (p<0.01), and Lefty2 (p<0.001) were also significantly decreased in iron-rich MEPs after week 3. Kif20b levels declined from week 2 to 3 (p<0.05) in iron-rich MEPs (B). n=3 per group.

Conversely, Lefty2 (0.2-fold) (p < 0.01), Ago1 (0.4-fold) (p < 0.01), Adra2a (0.48-fold) (p < 0.01) and Ldb1 (0.47-fold) (p < 0.01) were significantly reduced, after just 1 week of exposure to an iron-rich diet. Kif20b decreased significantly (0.38-fold) in iron-rich MEPs after 3 weeks, compared to week 2 (p < 0.05). Similarly, Ldb1 (0.74-fold) (p < 0.01), P2rx3 (0.17-fold) (p < 0.01), and Lefty2 (0.24-fold) (p < 0.001), decreased significantly in iron-rich MEPs on the 3^rd^ week. P2rx3 showed progressively declining expression from week 1 to week 3 in this cohort (Figure 7B).

#### 2.5. Investigating signalling pathways and downstream transcription factors which may regulate differential expression of megakaryopoiesis genes in MEPs

Finally, we assessed changes in probable signalling pathways and transcription factors, which may be responsible for upregulation of certain genes associated with megakaryopoiesis and platelet functions in iron-deficient MEPs. While Lefty2 is a member of the transforming growth factor-β (TGF-β) superfamily [13], other genes which showed elevated expression in iron-deficient MEPs, such as Ago1[14] and Kif20b [15] are closely associated with the TGF-β pathway. Past studies indicate platelets store and secrete TGF-β [16] and the latter transmit signals through Smad family of transcription factors [17].

We found that the gene encoding transforming growth factor-β receptor type 1(Tgfbr1) was increased by ∼ 12-fold in iron-deficient MEPs after 6 weeks, compared to MEPs isolated from mice fed on an iron-rich diet for 3 weeks (p < 0.01) (Figure 8A). Increased expression of Tgfbr1 directly resulted in significantly increased levels of R-Smads including Smad 2 by ∼ 12-fold in iron-deficient MEPs after 6 weeks (p < 0.05). Another R-Smad i.e. Smad 3 was also increased by more than 3-fold in iron-deficient MEPs after weeks 3 and 6, compared to MEPs isolated from mice fed on an iron-rich diet for just a week. (p < 0.05). (Figure 8B,C). This may also explain significantly increased expression of the Co-Smad i.e. Smad 4 in iron-deficient MEPs after 6 weeks (p < 0.05). (Figure 8D).

**Figure 8.**
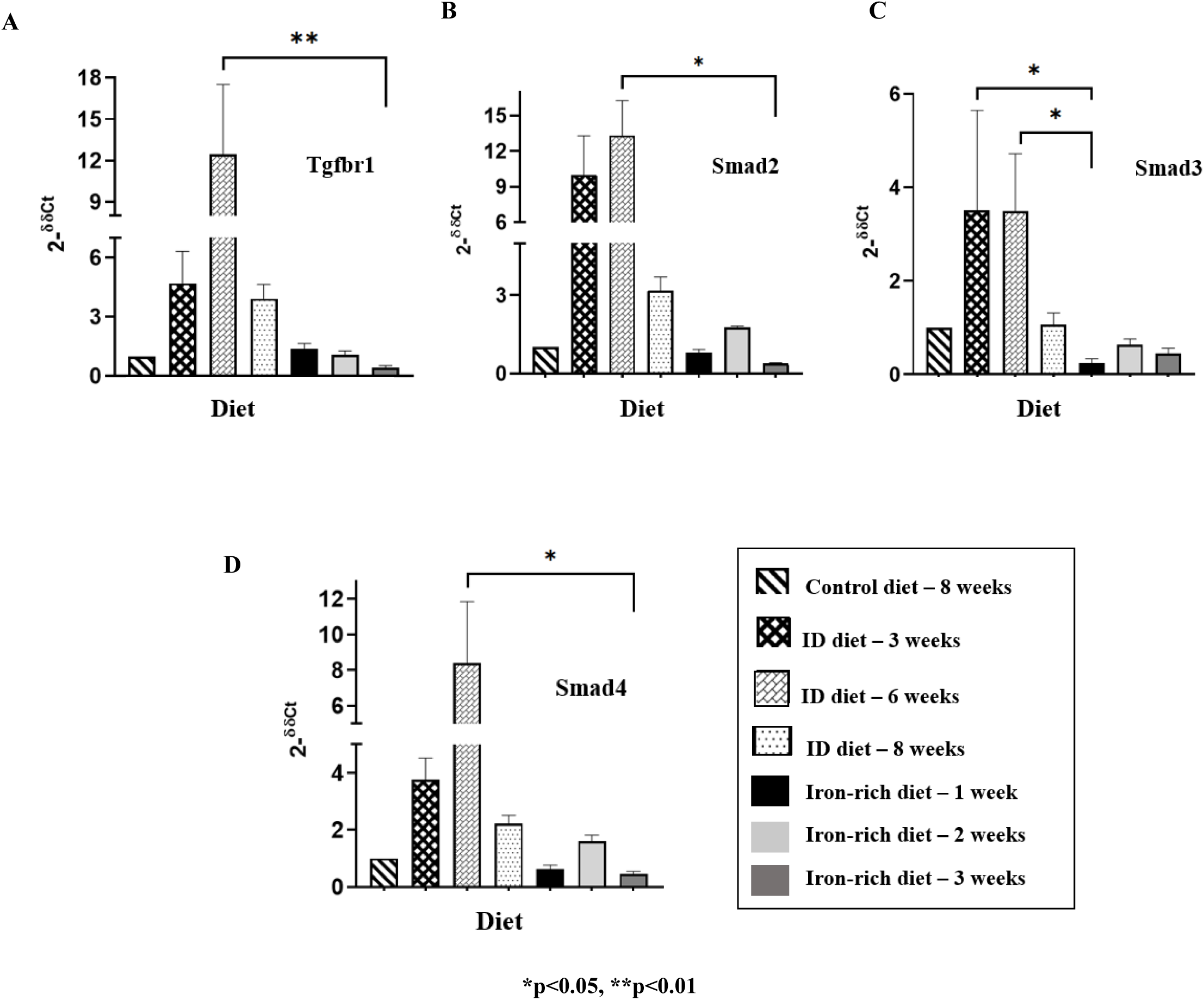
Exploring signalling pathways and downstream transcription factors regulating differential expression of megakaryopoiesis genes in MEPs –. Gene encoding transforming growth factor-β receptor type 1(Tgfbr1) showed increased expression in iron-deficient MEPs after 6 weeks (p<0.01), compared to iron-rich MEPs isolated on week 3 (A). Downstream transcription factors including R-Smads such as Smad-2 (B), Smad-3 (C) were increased in iron-deficient MEPs after 6 weeks (p<0.05), compared to iron-rich MEPs isolated after 3 weeks and 1 week respectively. Expression of the Co-Smad (Smad-4) was also increased in iron-deficient MEPs after 6 weeks (p < 0.05) (D). n=3 per group.

## 3. Discussion

The present study utilised male C57BL/6 mice to investigate the effects of varying iron levels in regulating megakaryopoiesis. We introduced a diet-based alteration in iron status in these mice by feeding them with control (40 ppm iron), iron-deficient (3 ppm) and iron-rich diets (20,000 ppm iron). Rishi et al. used a similar carbonyl iron-based diet to study hemochromatosis and hepcidin regulation in C57BL/6 mice [18]. We optimised the concentration of iron, which could induce iron deficiency, based on similar findings by Yook et al. [19]. We maintained mice on varying iron diets for different time durations to develop iron-deficient and iron-rich models. No stainable iron granules were observed in liver tissues in mice fed on an iron-deficient and control diet. However, mice maintained on an iron-rich diet developed Grade III iron overloading after 3 weeks. A recent study reported that 6-week-old C57BL/6 mice developed significant iron overloading while being maintained on a 2% carbonyl iron diet for 6 weeks [20]. The difference in results may be explained by the fact that younger mice, aged 4-5 weeks were used in our study, which developed iron overloading faster.

We further studied iron status by measuring the iron content of principal iron storage tissues [21]. Ippolito et al. recently reported that the hepatic and spleen iron content was significantly reduced in young C57BL/6 mice fed on an iron-deficient diet, containing less than 6 ppm of iron [22]. This was concordant with our findings, wherein the liver and spleen iron content were significantly higher in mice maintained on an iron-rich diet, than those exposed to an iron-deficient diet.

While assessing the effects of varying iron diets on haematological parameters, we observed that iron deficiency adversely affected Hb, RBC and MCV, which significantly declined after week 3. Riani et al. reported that young C57BL/6 mice maintained on an iron-deficient diet containing less than 6 ppm iron, had significantly reduced Hb. No significant differences were observed in other parameters[23]. Utilization of a lower concentration of iron in our study may explain why iron deficiency significantly affected haematological parameters. Like our findings, platelet counts were significantly elevated in iron-deficient mice in the study performed by Riani’s group. As our results closely reflected iron deficiency-associated thrombocytosis observed in IDA patients, we explored if iron deficiency affected the expression of megakaryocytic lineage-specific genes and transcription factors, in bone marrow cells.

Gata1 and Tal1 were significantly increased, when mice were maintained on an iron-deficient diet for 5 weeks. The critical role of Gata1 in megakaryocytic development has been reported in past *in vivo* studies. This may be due to the presence of functionally important Gata binding sites in most megakaryocytic promoter sequences [24]. Tal1 is essential for promoting megakaryocytic commitment. Its constitutive expression triggers a complex megakaryocytic transcriptional network. This, in turn, leads to enhanced *in vitro* production of megakaryocytes and platelets [25]. Interestingly, mice maintained on an iron-deficient diet also showed significantly increased expression of Mds and Evi1 complex locus (Mds1) and 3-phosphoinositide-dependent protein kinase 1 (Pdpk1). Mds1 acts as a transcriptional regulator, which is expressed in early precursor cells in the megakaryocytic lineage. Birdwell *et al.* have supported its role in promoting megakaryopoiesis in concert with other transcription factors [26]. Pdpk1 regulates F-actin cytoskeleton dynamics during megakaryocytic maturation and proplatelet formation [27]. The present study is the first one, to the best of our knowledge, which showed that certain well-known and recently reported megakaryocytic lineage-specific genes were significantly upregulated during iron deficiency in mice.

Recent evidence suggests that iron-dependent regulation of erythroid and megakaryocytic differentiation may extend beyond classical iron-responsive pathways. Heat shock cognate B (HSCB), a protein traditionally associated with iron-sulfur cluster delivery, acts as a critical regulator of both megakaryopoiesis and erythropoiesis by modulating activity of GATA1 and FOG1[28]. Thus, iron-dependent mechanisms may influence transcriptional networks governing lineage commitment, supporting the increased expression of megakaryocytic transcription factors observed in our study.

Results obtained from whole bone marrow cells, do not truly reflect the significance of iron in regulating megakaryopoiesis. To gain an understanding of the role of iron in regulating megakaryocytic commitment, megakaryocyte-erythroid progenitors (MEPs) i.e. Lin^-^ckit^+^Sca1^-^ CD34^-^ were isolated from bone marrow cells, and their gene expression profiles were analyzed by RNA Sequencing. The flow cytometric gating strategy used for identifying MEPs was similar to that employed by Zaro et al. [29].

An elegant study by Ferrucio *et al.* showed that low iron levels biased mice MEPs towards megakaryopoiesis. They developed a mouse model in which the gene encoding the “transmembrane serine protease 6” (TMPRSS6) was knocked out, which made them iron-deficient. These mice exhibited thrombocytosis. They reported that genes involved in regulating metabolic, vascular endothelial growth factor, and extracellular signal-regulated kinase (ERK) pathways were enriched in Tmprss6^−/−^ MEPs [11].

Our findings are further supported by a recent study demonstrating that iron availability regulates hematopoietic stem and progenitor cell (HSPC) fate decisions between erythroid and megakaryocytic lineages. While iron deficiency promoted thrombopoiesis, iron supplementation favoured erythropoiesis. Mechanistically, activation of the MAPK/ERK signalling pathway was identified as a key mediator of iron-regulated lineage commitment [30].

An interesting aspect of our study is that we induced an iron-deficient state in mice MEPs by a diet-based intervention without any genetic modifications. RNA isolated from MEPs from control, iron-deficient and iron-rich mice were subjected to RNA Sequencing. DEGs identified were filtered based on p values < 0.05. In iron-deficient MEPs, 148 genes were upregulated, and 156 genes were downregulated, compared to controls. We found that 138 genes were upregulated in iron-rich MEPs, while 231 genes were downregulated. Certain megakaryocytic transcription factors, including MYB, GATA1, TAL1, and GABPA were found in both iron-deficient and iron-rich MEPs (Supplementary figure 2).

We found relevant transcription factors and pathways associated with target DEGs from the Network Analyst and Reactome database, respectively. Genes involved in platelet degranulation, signalling, and homeostasis, such as Lefty2, Ptk2 and Col1a2, were differentially upregulated in iron-deficient MEPs. Other genes, including Ago1 and Kif20b, which regulate megakaryocytic development, were also upregulated in these MEPs. On the contrary, several genes associated with platelet homeostasis, aggregation, and megakaryocytic development, such as Gnb2, P2rx3, Adra2a and Ldb1 were differentially downregulated in iron-rich MEPs. These results pointed towards a crucial role of iron in influencing megakaryopoiesis in these progenitor stem cells.

These leads were further validated by gene expression analyses. We observed that levels of Lefty2, Gnb2, and P2rx3 increased significantly from week 3 to week 6 in iron-deficient MEPs. Most of these genes also showed increased expression after 8-9 weeks in iron-deficient MEPs. A contrasting scenario was observed in MEPs isolated from iron-rich mice. Lefty2, Ago1, Adra2a and Ldb1 levels were significantly reduced in iron-rich MEPs after a week. Kif20b, involved in regulation of megakaryocyte development and platelet production showed significant decline in expression over time in iron-rich MEPs. Most of the above-mentioned genes had significantly decreased levels in iron-rich MEPs after 3 weeks, compared to controls.

RNA sequencing results indicated that the gene encoding the Transforming growth factor β receptor1 (TGF-βR1) and Smad transcription factors (Smad 2, 3 and 4) were differentially expressed in iron-deficient and iron-rich MEPs. When we investigated the expression of these genes, we observed that both Tgfbr1 and Smad transcription factors were significantly increased in iron-deficient MEPs after 6 weeks. On the other hand, the iron status of MEPs affected the expression of genes regulating platelet signalling such as Lefty2, as well as those involved in megakaryocytic differentiation i.e. Ago1 and Kif20b. Interestingly, these genes have been closely linked with the TGF-β pathway [13],[14],[15]. Thus, iron deficiency may activate the TGF-β signalling pathway & consequently, downstream Smad transcription factors in MEPs. This in turn may influence lineage commitment towards megakaryopoiesis through elevated expression of different genes regulating megakaryocytic differentiation and platelet activation. These observations have been summarized in Figure 9.

**Figure 9.**
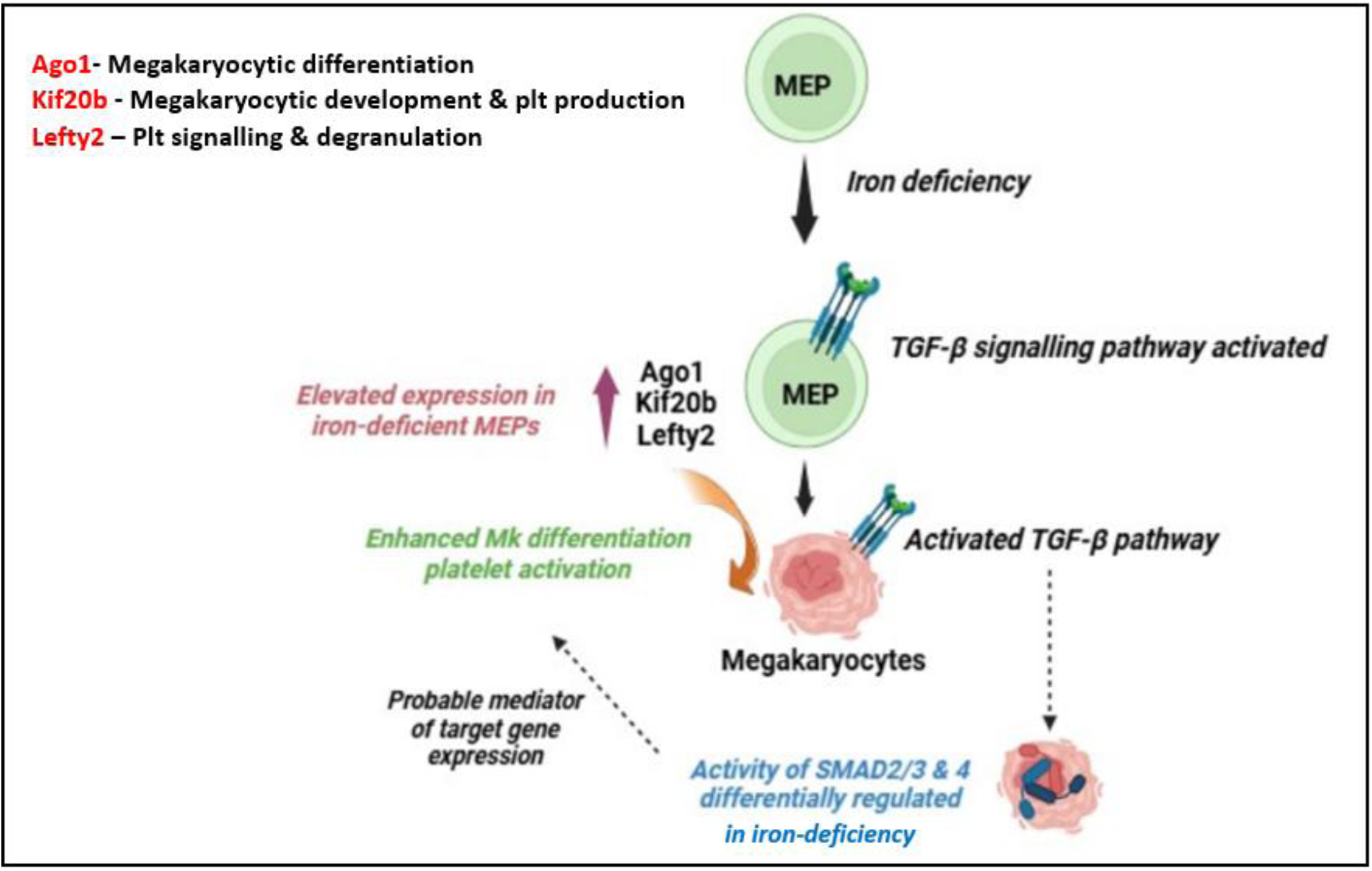
Iron deficiency is a key factor regulating megakaryocytic commitment in MEPs-. Our results indicate iron deficiency may cause differential regulation of the TGF-β signalling pathway and downstream Smad transcription factors in MEPs. This in turn may promote megakaryopoiesis, platelet production and activation through elevated expression of megakaryopoiesis related genes.

Apart from adult haematopoiesis, emerging evidence indicates that alterations in iron availability can influence developmental haematopoiesis and lineage specification. In a rat model of congenital iron deficiency, offspring born to iron-deficient dams exhibited significantly reduced hemoglobin levels, altered blood cell morphology, reduced platelet and white blood cell counts, despite postnatal iron recovery [31]. Concordant with our observations in adult MEPs, these studies underscore the broad regulatory role of iron in haematopoietic lineage determination.

## 4. Conclusion

The present study investigated the effects of varying iron levels in regulating megakaryopoiesis in an *in-viv*o context, by utilizing C57BL/6 mice maintained on different iron diets. An iron-deficient diet significantly decreased the hepatic iron content in these mice, while they had developed grade III iron-overloading, and significantly increased spleen and hepatic iron content on an iron-rich diet. Haematological parameters displayed a decreasing trend in iron-deficient mice, which also developed thrombocytosis.

Iron status affected the expression of different transcription factors and genes regulating megakaryopoiesis, such as Gata1, Tal1, Mds1 which showed significantly increased expression in bone marrow cells isolated from iron-deficient mice. To gain insights into the significance of iron levels in regulating the transcriptional switch towards megakaryopoiesis, MEPs were isolated from bone marrow cells of iron-deficient and iron-rich mice. An interesting aspect of our study is that we induced an iron-deficient and iron-rich state in MEPs solely by a diet-based intervention, which has been done by only a few past studies, to the best of our knowledge.

Several DEGs associated with megakaryocytic differentiation, platelet functions and homeostasis were upregulated in iron-deficient MEPs, while they were downregulated in iron-rich MEPs. Recent studies have further highlighted iron as an active regulator of haematopoietic lineage specification, and a bias of iron deficiency towards megakaryopoiesis, at the expense of erythropoiesis. These observations support our findings that iron deficiency induces transcriptional alterations in MEPs, leading to increased expression of genes associated with megakaryocyte differentiation and platelet production.

Among the upregulated DEGs, Lefty2, Ago1 and Kif20b have been closely associated with the TGF-β pathway. As genes encoding TGF-βR1 and Smad 2, 3 and 4 transcription factors were significantly increased in iron-deficient MEPs, we propose that iron deficiency may activate TGF-β signalling and downstream Smad-mediated transcriptional programs within MEPs. Thus, iron deficiency may act as a key upstream regulator of lineage-specifying molecular pathways and transcription factors, thereby promoting a shift in haematopoietic commitment towards megakaryopoiesis. These findings necessitate further validation by functional assays of identified target genes and transcription factors, to assess their contribution towards iron-dependent lineage commitment.

## Declarations

### Ethics approval statement

The present study was approved by the Institutional Animal Ethics Committee (IAEC) of CMC (IAEC No-5/2018) and the 2^nd^ Local Ethical Commission at the Warsaw University of Life Sciences, Poland (Permission No-WAW2/054/2020).

### Patient consent statement

NA

### Data Availability Statement

Data will be available upon request, by the corresponding author.

### Authors’ contribution statement

RD performed experiments and analyses, wrote, and edited the manuscript. LS helped in sample collection and processing. VM edited and reviewed the manuscript. SL supervised and AN, BK helped in statistical analyses of RNA Sequencing data. PL and RS supervised conductance of *in-vivo* experiments at IGHZ, Poland. ES designed the research proposal, analyzed the data, and reviewed the manuscript. All authors read and approved the final version of the manuscript.

