## Supplemental Figures and Table for "Investigating the significance of iron levels in influencing megakaryocytic commitment in megakaryocyte-erythroid progenitors"

**Supplementary Figure I- Mice models developed to study the role of iron in regulation of megakaryopoiesis by a diet-based intervention**

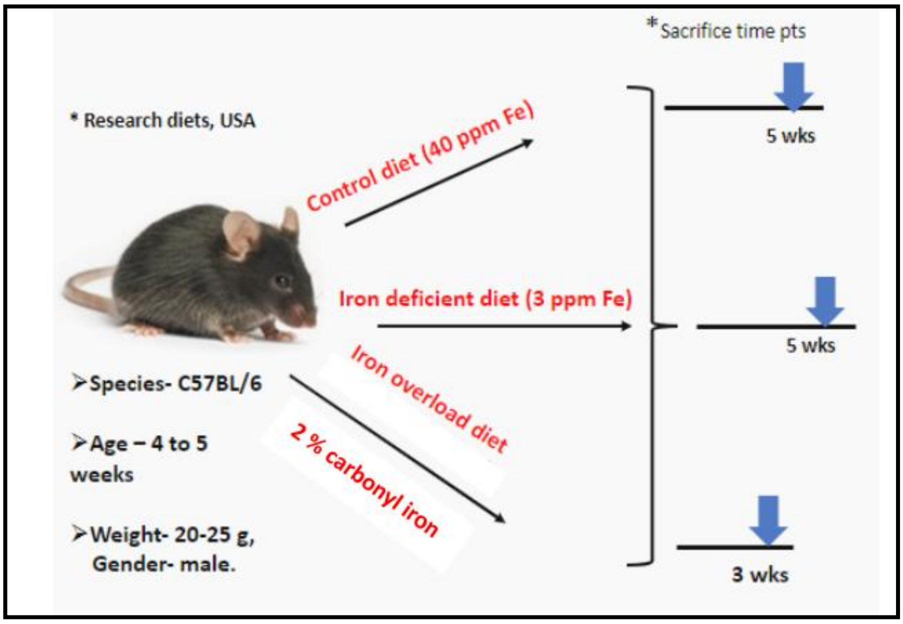


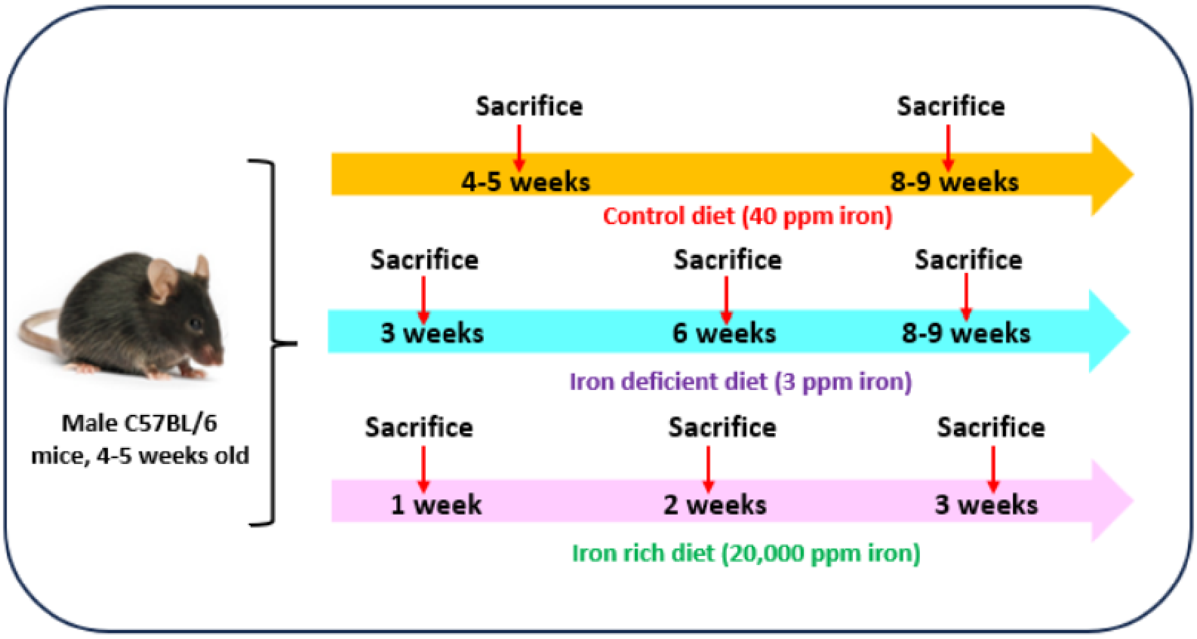

**Supplementary Figure 2 – Venn diagram of upregulated and downregulated DEGs in Iron-overloaded (IO), Iron-deficient (I) MEPs compared to controls (C)**

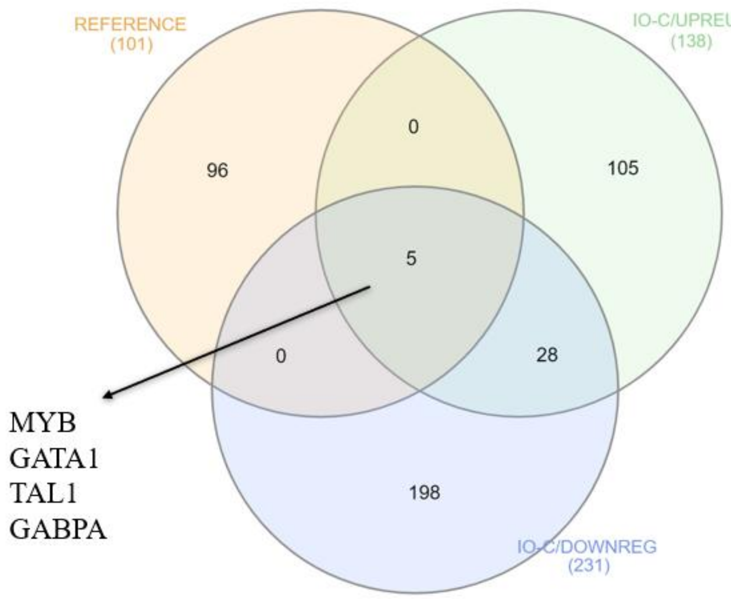


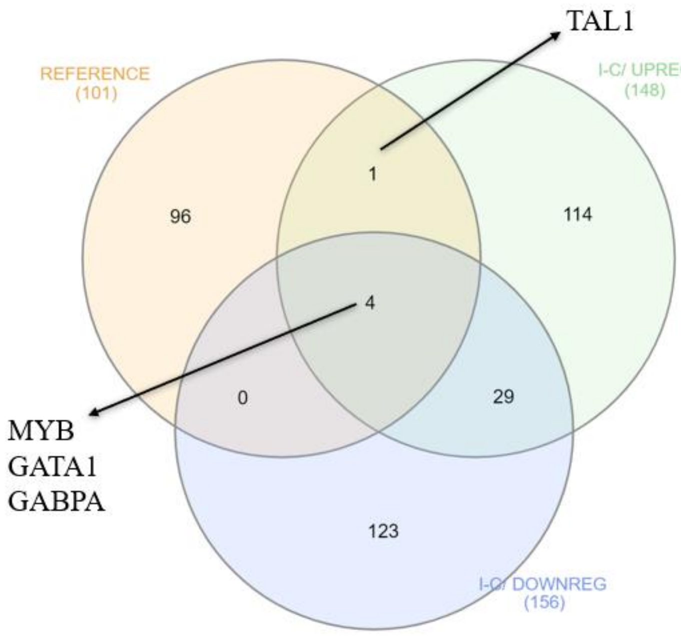

**Supplementary Table I- Primer sequences of megakaryocytic transcription factor encoding genes and genes associated with platelet activation and signalling**

| S.NO | Gene | Primer sequence |
|---|---|---|
| 1. | Gata1 | Forward 5'- GCTGAGTCCAGACCTCCTGA-3' |
| | | Reverse 5`- CACACTCTCTGGCCTCACAA-3' |
| 2. | Tal1 | Forward 5'- TCTGATGGTCCTCACACCAA-3' |
| | | Reverse 5`- GTGGGGATCAGCTTTCTGAG-3' |
| 3. | Mds1 | Forward 5'- ATCAGTGTCCCAAGGCATTT-3' |
| | | Reverse 5`- GCTAGGGTCCGTGAAAACCT-3' |
| 4. | Pdpk1 | Forward 5'- CGACGAAAAGCTGTATTTTGG-3' |
| | | Reverse 5`- CGTGTAAAACCGGGTACAGG-3' |
| 5. | Lefty2 | Forward 5'- CTCAAGGACTACGGAGCTCA -3' |
| | | Reverse 5`- CTAGGATCCAGTTCTCGGCC -3' |
| 6. | Gnb2 | Forward 5'- GATTCCATGTGCCGACAG -3' |
| | | Reverse 5`- GGTCAAAGAGGCGACAAG -3' |
| 7. | P2rx3 | Forward 5'- TTAAGATCGGCTGGGTGTGT -3' |
| | | Reverse 5`- CTGAAGTTGTAGCCAGGGGA -3' |
| 8. | Kif20b | Forward 5'- GCAAAGAAAGGGCTTATACTCGT -3' |
| | | Reverse 5`- CTATTGCCTTCCACAACTTCTGA -3' |
| 9. | Adra2a | Forward 5'- GTGACACTGACGCTGGTTTG -3' |
| | | Reverse 5`- CCAGTAACCCATAACCTCGTTG -3' |

| | | |
|---|---|---|
| 10. | Ago1 | Forward 5’- CAGCAGGTGTTTCAGGCAC -3’ |
| | | Reverse 5`- GGACACTTATCCGGCTTGATG -3’ |
| 11. | Ldb1 | Forward 5’- CTGGACAGAGGAGTGTGACA -3’ |
| | | Reverse 5`- GGTCCATCCTCCAAGCAGAA -3’ |
| 12. | Gapdh | Forward 5’- CTCCCACTCTTCCACCTTCG -3’ |
| | | Reverse 5`- GCCTCTCTTGCTCAGTGTCC -3’ |
